# Electrophilic PROTACs that degrade nuclear proteins by engaging DCAF16

**DOI:** 10.1101/443804

**Authors:** Xiaoyu Zhang, Vincent M. Crowley, Thomas G. Wucherpfennig, Melissa M. Dix, Benjamin F. Cravatt

## Abstract

Ligand-dependent protein degradation has emerged as a compelling strategy to pharmacologically control the protein content of cells. So far, only a limited number of E3 ligases have been found to support this process. Here, we use a chemical proteomic strategy to discover that DCAF16 – a poorly characterized substrate recognition component of CUL4-DDB1 E3 ubiquitin ligases – promotes nuclear-restricted protein degradation upon modification by cysteine-directed heterobifunctional electrophilic compounds.

---

Conventional small-molecule probes and drugs act by directly perturbing the functions of proteins (e.g., blocking enzyme catalysis or antagonizing receptor signaling). Many proteins, however, possess multiple functional domains, and therefore a compound that binds to only one of these domains may fail to fully inactivate the protein. An alternative emerging strategy uses chemical probes that direct proteins to the proteolytic degradation machinery of the cell, leading to the loss of protein expression^1, 2^. This targeted protein degradation approach leverages two types of small molecules – those that form tripartite complexes with specific E3 ubiquitin ligases and neosubstrate proteins (e.g., the IMiD class of therapeutics^3, 4^) and bifunctional compounds, often referred to as PROTACs (proteolysis targeting chimeras), which couple E3 ligase ligands to substrate ligands via a variably structured linker^5-7^. Targeted protein degradation has the potential to act in a catalytic manner^8^ that may lower the drug concentrations required to produce a pharmacological effect and operate sub-stoichiometrically to avoid antagonizing the natural functions of the participating E3 ligase. Despite these advantages, to date, only a handful of the 600+ human E3 ligases have been found to support ligand-mediated protein degradation^5-7, 9-11^. Importantly, these E3 ligases has been found to show distinct and restricted substrate specificities^12, 13^, a parameter that is not yet predictable or easily controlled, underscoring the need to discover additional ligandable E3 ligases with differentiated properties to realize the full scope of targeted protein degradation as a pharmacological strategy.

We have recently introduced chemical proteomic platforms for globally and site-specifically mapping the reactivity of electrophilic small molecules in native biological systems^14-16^. These experiments have identified a subset of fragment electrophiles, referred to hereafter as ‘scouts’^15^, that capture a remarkably broad fraction of the 100s-1000s of covalent small molecule-cysteine interactions discovered so far by chemical proteomics. We surmised herein that these scout fragments, when incorporated into heterotypic bifunctional compounds containing optimized protein ligands, may offer an expedited path to discover E3 ligases capable of supporting targeted protein degradation through covalent reactivity with cysteine residues. With this goal in mind, we fused three scout fragments – KB02, KB03, and KB05 – to the SLF ligand that binds tightly and selectively to FKBP12 (**Fig. 1a**), a cytosolic prolyl isomerase that has been frequently used to study ligand-induced protein degradation^17, 18^. We then stably expressed FLAG-tagged variants of FKBP12 (FLAG-FKBP12) or FKBP12 with a *C*-terminal nuclear localization sequence (FLAG-FKBP12_NLS) in HEK293T cells to provide cell models for evaluating cytosolic-and nuclear-localized E3-mediated degradation pathways, respectively (**Fig. 1b** and **Supplementary Fig. 1a**).

**Figure 1.**
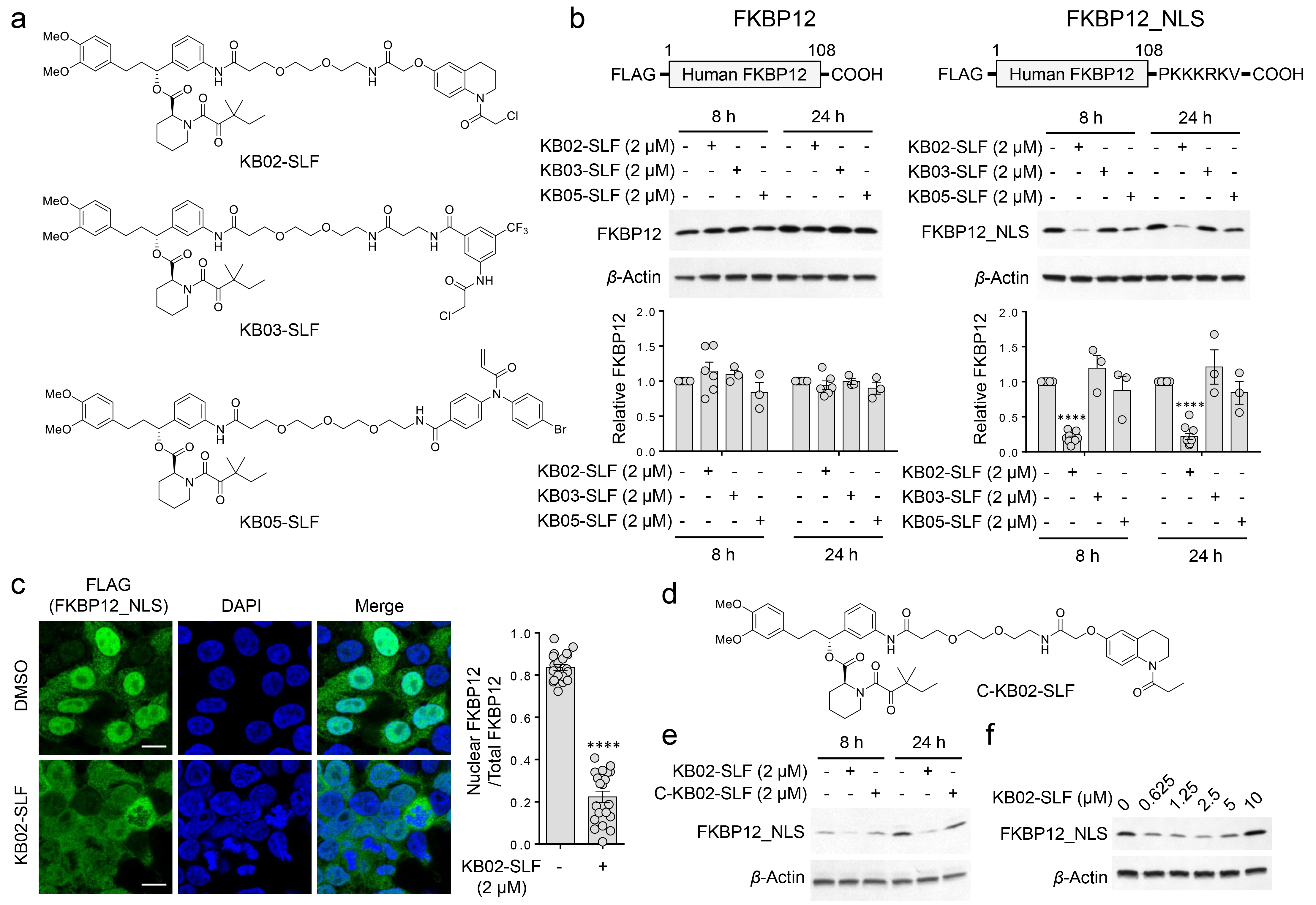
Discovery of an electrophilic PROTAC that degrades nuclear FKBP12. **a**, Structures of electrophilic bifunctional compounds, or PROTACS, containing scout fragments (KB02, KB03, and KB05) coupled to the FKBP12 ligand SLF. **b**, Western blot using anti-FLAG antibody of cytosolic (FLAG-FKBP12) and nuclear (FLAG-FKBP12_NLS) FKBP12 proteins expressed by stable transduction in HEK293T cells following 8 or 24 h of treatment with electrophilic PROTACs (2 μM). Bar graph represents quantification of the relative FKBP12 protein content, with DMSO-treated cells set to a value of 1. Data represent mean values ± SEM for 3-10 biological replicates. Statistical significance was calculated with unpaired two-tailed Student’s t-tests comparing DMSO- to KB02-SLF-treated samples. \*\*\*\**P* < 0.0001. **c**, Immunofluorescence using anti-FLAG antibody of FLAG-FKBP12_NLS in HEK293T cells following treatment with DMSO or KB02-SLF (2 μM, 8 h). Bar graph (right) represents quantification of the relative nuclear to whole cell immunostaining for DMSO- and KB02-SLF-treated samples. Data represent mean values ± SEM (n = 20 from two biological replicates). Statistical significance was calculated with unpaired two-tailed Student’s t-tests comparing DMSO- to KB02-SLF-treated samples. \*\*\*\**P* < 0.0001. Scale bar, 10 μm. **d**, Structure of non-electrophilic control compound (C-KB02-SLF). **e**, Western blot of FLAG-FKBP12_NLS in HEK293T cells treated with KB02-SLF or C-KB02-SLF (2 μM, 8 or 24 h). The result is a representative of three independent experiments. **f**, Concentration-dependent degradation of FLAG-FKBP12_NLS by KB02-SLF in HEK293T cells (24 h treatment with indicated concentrations of KB02-SLF). The result is a representative of four independent experiments.

We first confirmed by anti-FLAG western blotting that both cytosolic and nuclear FKBP12 are degraded by lenalidomide-SLF (**Supplementary Fig. 1b, c**), a bifunctional molecule comprised of SLF coupled to lenalidomide, a ligand for the E3 ligase cereblon (CRBN)^17, 18^. We next evaluated the electrophilic scout fragment-SLF bifunctional compounds for effects on FKBP12 degradation in HEK293T cells. Under the treatment conditions (2 μM, 8 or 24 h), none of the compounds altered cytosolic FKBP12 (**Fig. 1b**). On the other hand, the KB02-SLF compound promoted a substantial reduction in nuclear FKBP12 (**Fig. 1b**) that was sustained across a 4-72 h time frame (**Supplementary Fig. 2a**). Cell imaging studies confirmed the selective loss of nuclear-localized FKBP12 in KB02-SLF-treated cells (**Fig. 1c**). These imaging studies, along with western blotting experiments (**Supplementary Fig. 1a**), pointed to a fraction of FKBP12 that remained cytosolically localized in FLAG-FKBP12_NLS-transfected cells and was consequently unaffected by KB02-SLF treatment (**Fig. 1c**).

Reductions in FKBP12_NLS were not observed with an analogue of KB02-SLF where the electrophilic alpha-chloroacetamide group was replaced with an unreactive propanamide (C-KB02-SLF; **Fig. 1d, e**), indicating that the mechanism of action of KB02-SLF involved covalent modification of one or more proteins. KB02-SLF, as well as analogues of this compound with different linker lengths (**Supplementary Fig. 2b**), promoted the loss of FKBP12_NLS across a concentration of ~0.5-5 μM, but showed varied reductions in activity at higher concentrations (**Fig. 1f** and **Supplementary Fig. 2b**). Such parabolic concentration-dependence is a feature of PROTACs, where binary complexes gain prevalence over ternary complexes at higher cellular concentrations of compound^19^. Consistent with this mode of action, we did not observed reductions in FKBP12_NLS in cells treated with the two separate components (KB02 and SLF) of the KB02-SLF bifunctional compound (**Supplementary Fig. 2c**). Pre-treatment with SLF did, on the other hand, block the KB02-SLF-induced loss of FKBP12_NLS (**Supplementary Fig. 2d**). We also found that KB02-SLF induced polyubiquitination of nuclear FKBP12_NLS (**Fig. 2a** and **Supplementary Fig. 3**), but not cytosolic FKBP12 (**Fig. 2a**) and KB02-SLF-mediated loss of FKBP12_NLS was blocked by the proteasome inhibitor MG132 (**Fig. 2b**) and neddylation inhibitor MLN4924^20^ (**Fig. 2c**). Similar effects of KB02-SLF on FKBP12_NLS were observed in a second human cell line (MDA-MB-231) (**Supplementary Fig. 4**). These data, taken together, supported that KB02-SLF promoted the proteasomal degradation of nuclear-localized FKBP12 via the action of a Cullin-RING ubiquitin ligase (CRL).

**Figure 2.**
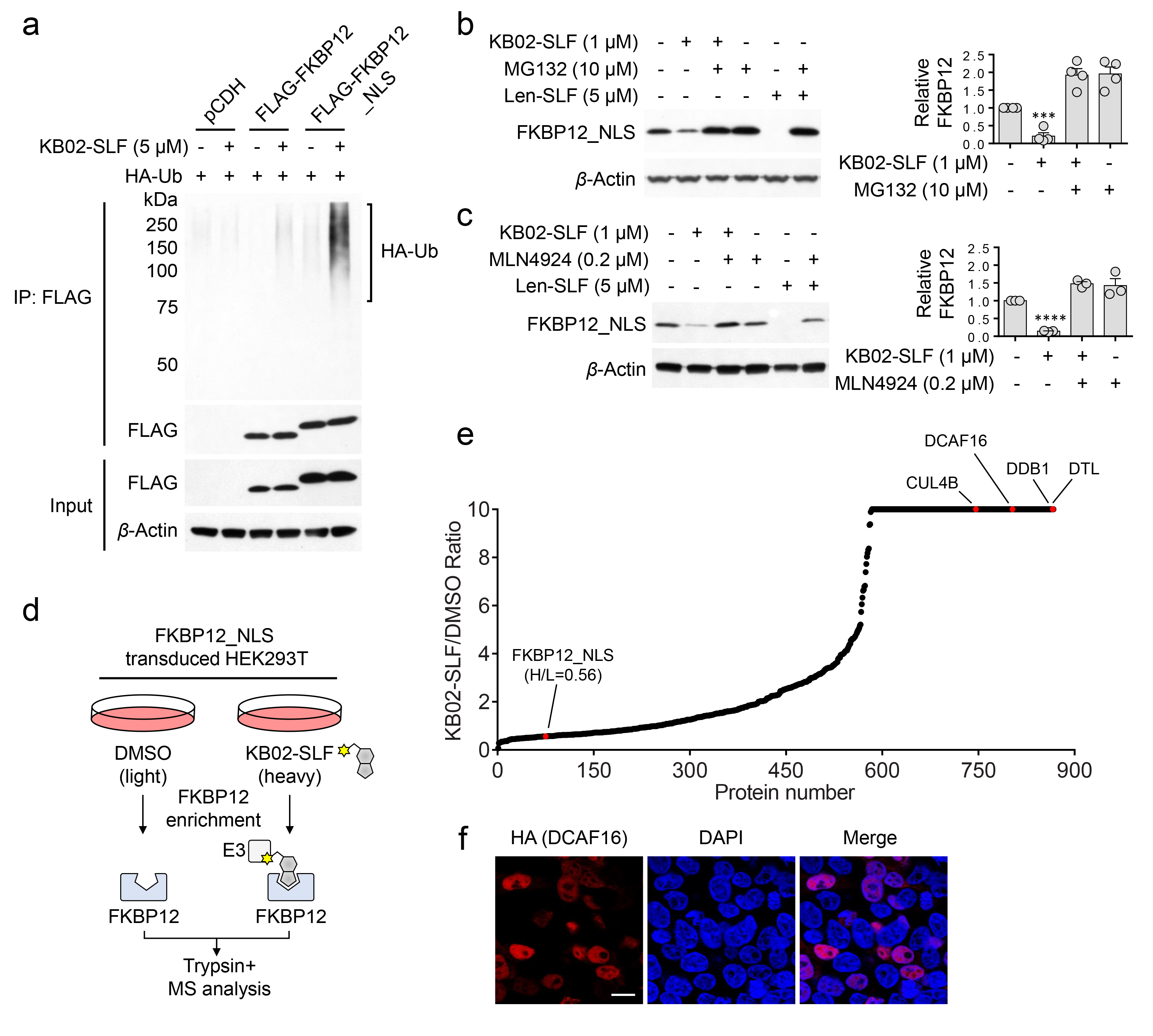
Characterization of KB02-SLF-mediated degradation of nuclear FKBP12. **a**, KB02-SLF mediates polyubiquitination of nuclear (FLAG-FKBP12_NLS), but not cytosolic (FLAG-FKBP12) in HEK293T cells. HEK293T cells stably expressing FLAG-FKBP12 or FLAG-FKBP12_NLS were transiently transfected with HA-Ubiquitin (HA-Ub) for 24 h and then treated with DMSO or KB02-SLF (5 μM) in the presence of the proteasome inhibitor MG132 (10 μM) for 2 h. Anti-FLAG-immunoprecipitated FLAG-FKBP12 and FLAG-FKBP12_NLS proteins were analyzed for ubiquitination by Western blotting using an anti-HA antibody. The result is a representative of three independent experiments. **b**, **c**, KB02-SLF-mediated FLAG-FKBP12_NLS degradation is blocked by MG132 (**b**) and the neddylation inhibitor MLN4924 (**c**). HEK293T cells stably expressing FLAG-FKBP12_NLS were co-treated with KB02-SLF (1 μM) and MG132 (10 μM) or MLN4924 (1 μM) for 8 h. Len-SLF is a PROTAC compound comprised of lenalidomide coupled to SLF and was used as a positive control. Bar graph (right) represents quantification of the relative FKBP12 protein, with DMSO-treated cells set to a value of 1. Data represent mean values ± SEM for 3-4 biological replicates. Statistical significance was calculated with unpaired two-tailed Student’s t-tests comparing DMSO- to KB02-SLF-treated samples with or without MG132 or MLN4924. \*\*\**P* < 0.001; \*\*\*\**P* < 0.0001. **d**, Schematic for identifying KB02-SLF-recruited E3 ubiquitin ligases by anti-FLAG affinity enrichment coupled to mass spectrometry (MS)-based proteomics. Light and heavy amino acid-labeled HEK293T cells stably expressing FLAG-FKBP12_NLS were treated with DMSO or KB02-SLF (10 μM), respectively, for 2 h in the presence of MG132 (10 μM). Light and heavy cells were then lysed, subject to anti-FLAG immunoprecipitation, and the affinity-enriched proteins combined, digested with trypsin, and analyzed by LC-MS/MS. **e**, SILAC heavy/light (H/L) ratio values of proteins identified in anti-FLAG affinity enrichment experiments (outlined in part **d**), where a high ratio indicates proteins selectively enriched from cells treated with KB02-SLF. The result is a representative of three independent experiments. **f**, Immunofluorescence using an anti-HA antibody showing nuclear localization of HA-DCAF16 (expressed by transient transfection in HEK293T cells). Scale bar, 10 μm. The result is a representative of two independent experiments.

Working under a model where KB02-SLF induced the degradation of FKBP12_NLS by covalently reacting with one or more CRL(s), we first attempted to identify the recruited CRL(s) using a chemical proteomic method termed isoTOP-ABPP (isotopic tandem orthogonal proteolysis-activity-based protein profiling), where ligand-engaged cysteines in cells are mapped by competitive blockade of reactivity with an iodoacetamide (IA)-alkyne probe^21^. However, we did not observe any cysteines on CRL(s) that were strongly engaged (> 75%) by KB02-SLF (10 μM, 2 h) in HEK293T cells by isoTOP-ABPP (**Supplementary Table 1**). We alternatively considered that KB02-SLF may show low stoichiometry engagement of a CRL, which would then function catalytically to drive FKBP12_NLS degradation. Such low fractional occupancy ligand-protein interactions are not easily mapped by competitive profiling methods like isoTOP-ABPP, as they would minimally perturb IA-alkyne-cysteine reactions. We therefore employed an alternative proteomic approach, in which FLAG-mediated affinity enrichment was used to identify proteins that associated with FKBP12_NLS in a KB02-SLF-dependent manner (**Fig. 2d**). This experimental set up identified two substrate receptor components of CRLs – DCAF16 and DTL – that were substantially enriched (> 5-fold) in HEK293T cells treated with KB02-SLF compared to DMSO-treated control cells (**Fig. 2e**, **Supplementary Fig. 5a** and **Supplementary Table 2**). Neither DCAF16 nor DTL were enriched by KB02-SLF in an additional control experiment performed with mock-transfected cells lacking FKBP12_NLS (**Supplementary Fig. 5a, b** and **Supplementary Table 2**). We also observed KB02-SLF-dependent enrichment of DCAF16 in MDA-MB-231 cells (**Supplementary Fig. 4d, e** and **Supplementary Table 2**).

Notably, both DCAF16 and DTL are predicted to be nuclear proteins (https://psort.hgc.jp/). DTL (also known as CDT2 and DCAF2) plays a key role in cell cycle control and DNA damage response^22^. DCAF16, on the other hand, remains poorly characterized and shares negligible homology with other DDB1-and CUL4-associated factor (DCAF) proteins that participate in CRLs^23^. We confirmed the nuclear localization of an HA-tagged DCAF16 in transfected HEK293T cells (**Fig. 2f**). Consistent with KB02-SLF engaging functional CRL complexes potentially containing DCAF16 and/or DTL, we also observed enrichment of additional CRL components – DDB1 and CUL4B – in KB02-SLF-treated cells (**Fig. 2e** and **Supplementary Fig. 5a**). We next used small hairpin RNA (shRNA)-mediated knockdown to find that reductions in DCAF16, but not DTL, substantially prevented KB02-SLF-mediated degradation of FKBP12_NLS (**Supplementary Fig. 5c, d**). shRNA-knockdown of DCAF16 also blocked KB02-SLF-induced polyubiquitination of FKBP12_NLS (**Supplementary Fig. 5e**). We further verified the involvement of DCAF16 in KB02-SLF-mediated degradation of FKBP12_NLS by CRISPR/Cas9 genetic knockout^24^. We used genomic sequencing and mass spectrometry-based proteomics to confirm the genetic disruption of DCAF16 in three independent clones (DCAF16-/-) compared to three wild type DCAF16 clones (DCAF16+/+) (**Supplementary Fig. 6a-c** and **Supplementary Table 3**). KB02-SLF supported the degradation of FKBP12_NLS in all three DCAF16+/+ clones, but not in any of the three DCAF16-/- clones (**Fig. 3a** and **Supplementary Fig. 6d**). Moreover, recombinant expression of HA-tagged DCAF16 restored KB02-SLF-induced FKBP12_NLS degradation in DCAF16-/- cells (**Fig. 3b**) and further enhanced the extent of KB02-SLF-induced FKBP12_NLS degradation in DCAF16+/+ cells (**Supplementary Fig. 6e**). We did not observe the degradation of HA-DCAF16 itself following treatment of cells with KB02-SLF (**Fig. 3b** and **Supplementary Fig. 6e, f**). These data, taken together, demonstrate that the degradation of FKBP12_NLS induced by KB02-SLF is mediated by DCAF16.

**Figure 3.**
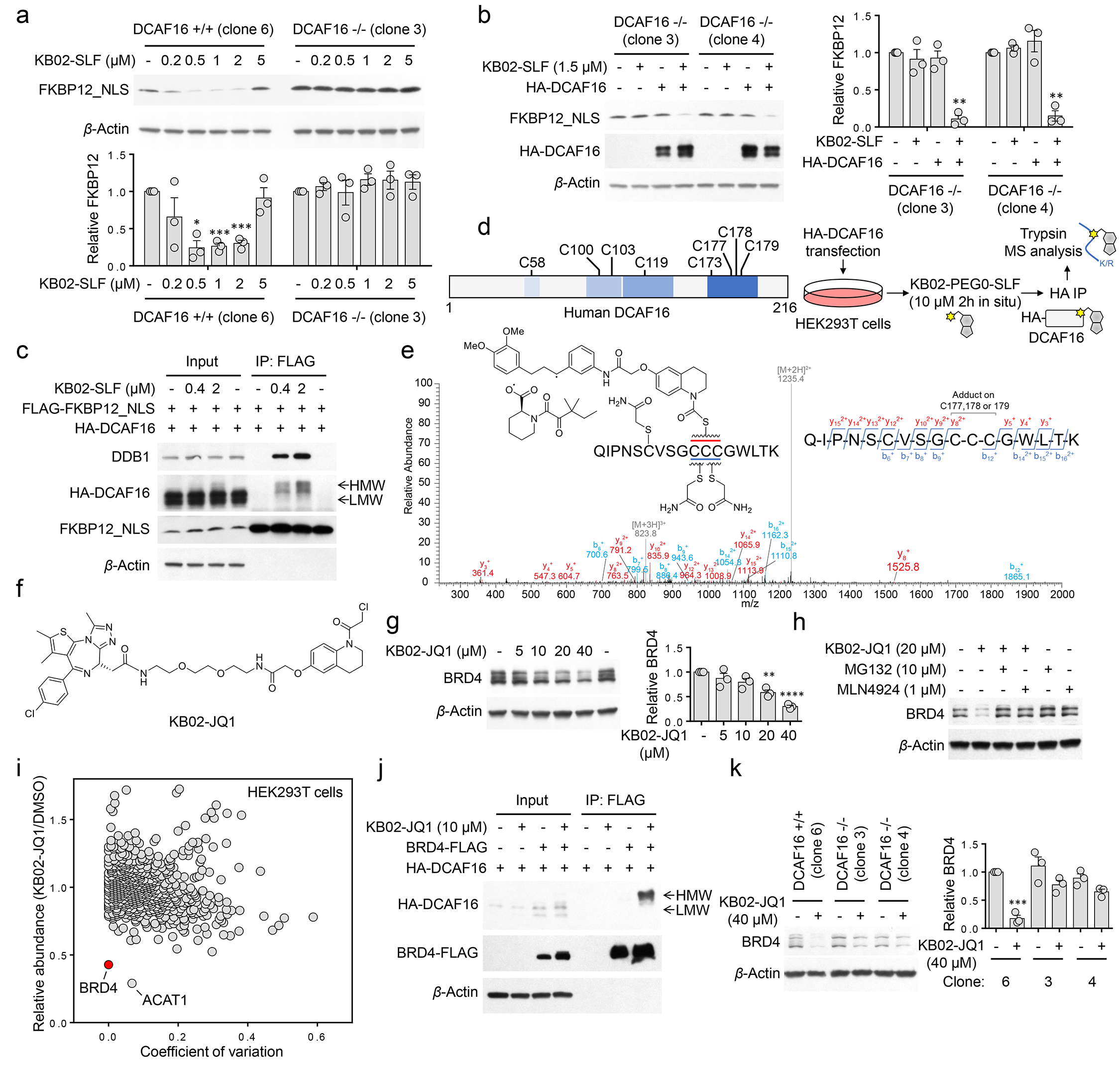
DCAF16 mediates electrophilic PROTAC-induced degradation of nuclear proteins. **a**, Concentration-dependent degradation of stably expressed FLAG-FKBP12_NLS in DCAF16+/+ (clone 6) and DCAF16-/- (clone 3) HEK293 cells following treatment with KB02-SLF for 8 h. Bar graph (bottom) represents quantification of the relative FKBP12 protein content, with DMSO-treated cells set to a value of 1. Data represent mean values ± SEM for 3 biological replicates. Statistical significance was calculated with unpaired two-tailed Student’s t-tests comparing DMSO- to KB02-SLF-treated samples. \**P* < 0.05; \*\*\**P* < 0.001. **b**, Expression of HA-DCAF16 in DCAF16-/- cells restored KB02-SLF-mediated degradation of FLAG-FKBP12_NLS. DCAF16-/- cells were transiently transfected with FLAG-FKBP12_NLS and either HA-DCAF16 or empty pRK5 vector as a control for 24 h and then treated with KB02-SLF (1.5 μM, 8 h). Bar graph (right) represents quantification of the relative FKBP12 protein content, with DMSO-treated cells set to a value of 1. Data represent mean values ± SEM for 3 biological replicates. Statistical significance was calculated with unpaired two-tailed Student’s t-tests comparing DMSO- to KB02-SLF-treated samples. \*\**P* < 0.01. **c**, A higher molecular weight (HMW) form of HA-DCAF16 is observed in HEK293T cells treated with KB02-SLF (0.4 or 2 μM, 2 h in the presence of 10 μM MG132), and this HMW form, but not the lower molecular weight (LMW) form of HA-DCAF16 co-immunoprecipitated with FLAG-FKBP12_NLS. The result is a representative of three independent experiments. **d**, Left: schematic representation of human DCAF16 protein denoting eight cysteines. Right: schematic for identifying KB02-PEG0-SLF modified cysteines on DCAF16. **e**, MS/MS spectrum of KB02-PEG0-SLF-modified, triply charged DCAF16 peptide (amino acids 168-184; see **Supplementary Fig. 8a**). The *m/z* 1235.4 ion represents a doubly charged DCAF16 peptide (168-184) with a cleaved probe adduct that occurs in MS/MS. The b- and y-ions are shown along with the peptide sequence. **f**, Structure of KB02-JQ1. **g**, Concentration-dependent degradation of endogenous BRD4 in HEK293T cells following treatment with KB02-JQ1 for 24 h. Bar graph (right) is quantification of the relative BRD4 level. The BRD4 level from DMSO-treated cells is set to 1. Data represent mean values ± SEM for 3 biological replicates. Statistical significance was calculated with unpaired two-tailed Student’s t-tests comparing DMSO- to KB02-JQ1-treated samples. \*\**P* < 0.01; \*\*\*\**P* < 0.0001. **h**, KB02-JQ1-mediated BRD4 degradation is blocked by proteasome inhibitor MG132 and neddylation inhibitor MLN4924. HEK293T cells were preincubated with 10 μM MG132 or 1 μM MLN4924 for 4 h, followed by 20 h of treatment with 20 μM KB02-JQ1 and 10 μM MG132 or 1 μM MLN4924. The result is a representative of three independent experiments. **i**, Fold-change in protein abundance between heavy- and light-isotopically labeled HEK293T cells treated with KB02-JQ1 (20 μM) or DMSO, respectively, for 24 h. The y-axis and x-axis correspond to the average relative abundance (DMSO/KB02-JQ1) and coefficient of variation, respectively, from duplicate experiments. The average relative abundance of each protein is normalized to the average abundance of proteins in control DMSO/DMSO samples (to account for slight deviations in heavy isotope incorporation; **Supplementary Table 3**) and represents mean values from two independent experiments. **j**, FLAG tagged BRD4 co-immunoprecipitated with HA-DCAF16 in the presence of KB02-JQ1. HEK293T cells were co-transfected with BRD4-FLAG and HA-DCAF16 or pRK5 vector for 24 h and treated with 10 μM KB02-JQ1 or DMSO and 10 μM MG132 for 2 h. **k**, Degradation of BRD4 in HEK293 DCAF16+/+ (clone 6) and -/- (clone 3 and 4) cells following 24 h of treatment with 40 μM KB02-JQ1. Bar graph (right) represents quantification of the relative BRD4 protein content, with DMSO-treated cells set to a value of 1. Data represent mean values ± SEM for 3 biological replicates. Statistical significance was calculated with unpaired two-tailed Student’s t-tests comparing DMSO- to KB02-JQ1-treated samples. \*\*\**P* < 0.001.

In further support of ternary complex formation involving a DCAF16-CRL, we found that HA-DCAF16 and DDB1 co-immunoprecipitated with FLAG-FKBP12_NLS in the presence of KB02-SLF (**Fig. 3c**) or its linker analogues (**Supplementary Fig. 7a**). In the course of performing these co-immunoprecipitation experiments, we noticed that KB02-SLF-treated cells showed a higher molecular weight (HMW) form of HA-DCAF16, consistent with covalent modification of this protein by KB02-SLF (**Fig. 3c** and **Supplementary Fig. 7a-f**). Only a modest fraction of HA-DCAF16 was converted to this HMW in the presence of increasing concentrations of KB02-SLF (**Fig. 3c** and **Supplementary Fig. 7b**). Notably, however, co-immunoprecipitations with FKBP12_NLS exclusively pulled down the HMW, but not LMW form of DCAF16 (**Fig. 3c** and **Supplementary Fig. 7**), supporting a ternary complex model where KB02-SLF is covalently and non-covalently bound to DCAF16 and FKBP12_NLS, respectively. Also consistent with this model, DCAF16 was not co-immunoprecipitated from cells treated with the non-electrophilic control compound C-KB02-SLF (**Supplementary Fig. 7c**) and minimally enriched from cells treated with the other two electrophilic bifunctional compounds (KB03-SLF and KB05-SLF) that did not support FKBP12_NLS degradation (**Supplementary Fig. 7d, e**).

DCAF16 is a 216 aa protein that is highly conserved across mammals (e.g., human and rabbit DCAF16 share 97% identity), but absent in rodents. The DCAF16 protein has eight cysteine residues, including a cluster of four cysteines between amino acids (aa) 173-179 (**Fig. 3d**). We next attempted to map KB02-PEG0-SLF-reactive cysteine(s) in DCAF16 by treating HA-DCAF16-transfected HEK293T cells with DMSO or KB02-PEG0-SLF (10 μM, 2 h), affinity purifying HA-DCAF16, and subjecting the protein to trypsin digestion and LC-MS/MS analysis (**Fig. 3d**). We searched the MS1 (parent ion) profiles for the *m/z* values of unmodified and KB02-PEG0-SLF-modified DCAF16 tryptic peptides and identified a KB02-PEG0-SLF-modified form of the tryptic peptide (aa 168-184) containing C173 and C177-179 (**Supplementary Fig. 8a**). In contrast, we did not identify KB02-PEG0-SLF-modified forms for tryptic peptides containing C100/C103 (aa 97-106) or C119 (aa 107-133) (**Supplementary Fig. 8b, c**). We did not detect the tryptic peptide containing C58, likely because it is a very small in length (four aa). Tandem (MS/MS) analysis of the KB02-PEG0-SLF-modified DCAF16 peptide indicated that the most likely modified residue(s) was C177, C178, and/or C179 (**Fig. 3e**). Our efforts to evaluate the contribution of C177-179 and other cysteine residues to KB02-SLF-induced degradation of FKBP12_NLS by site-directed mutagenesis were hindered, in part, by the dramatic reduction in expression of cysteine-to-serine mutants for multiple residues (e.g., C177, C179) (**Supplementary Fig. 9a**). These mutagenesis studies did reveal that the C58S-, C173S-, and C178S-DCAF16 mutants were expressed at near-wild type levels and supported KB02-SLF-induced degradation of FKBP12_NLS (**Supplementary Fig. 9a, b**), indicating that C58, C173, and C178 are unlikely to constitute primary sites of engagement for KB02-SLF. We additionally analyzed whether cysteine-to-serine mutants of DCAF16 could form a ternary complex with KB02-SLF and FKBP12_NLS in cells pre-treated with the proteasome inhibitor MG132, which we found to normalize DCAF mutant protein expression levels (**Supplementary Fig. 9c**). Among the five mutants tested, C58S-, C173S- and C178S-DCAF16, but not C177S- and C179S-DCAF16, co-immunoprecipitated with FKBP12_NLS in KB02-SLF-treated cells (**Supplementary Fig. 9c**). While our data, taken in total, support C177 and/or C179 in DCAF16 as possible sites of modification responsible for KB02-SLF-induced degradation of FKBP12_NLS, we should qualify this interpretation by noting the potential for improper folding of the C177S- and C179S-DCAF16 mutants as a confounding factor.

We finally investigated whether covalent modification of DCAF16 could support the degradation of another nuclear protein. For these studies, we selected BRD4 as the nuclear protein, as it has a potent and selective ligand JQ1 that has been successfully coupled to other E3 ligands to promote degradation^7, 17, 25, 26^. We found that a KB02-JQ1 bifunctional compound (**Fig. 3f**) promoted, in a concentration-dependent manner, the degradation of BRD4 in HEK293T cells (**Fig. 3g**), and this effect was blocked by MG132 or MLN4924 (**Fig. 3h**). BRD4 was not degraded in cells treated with either the two separate components (KB02 and JQ1) of KB02- JQ1 or the FKBP12-directed bifunctional compound KB02-SLF (**Supplementary Fig. 10a**). We noted that much higher concentrations of KB02-JQ1 (20-40 μM) were required to degrade BRD4 compared to the degradation of FKBP12_NLS by KB02-SLF (0.5-2 μM). We suspect that this apparent difference in potency may reflect differential cellular uptake of the two KB02 bifunctional compounds, as KB02-JQ1 also showed a rightward shift in cytotoxicity (IC_50_ > 50 μM) compared to KB02-SLF and the parent electrophilic fragment KB02 (IC_50_ values of 14-23 μM) (**Supplementary Fig. 10b**). We then assessed the proteome-wide effect of KB02-JQ1 treatment in HEK293T cells and found that the KB02-JQ1-induced degradation of BRD4 was selective, with only one additional protein – ACAT1 – showing similar reductions in KB02-JQ1-treated cells (**Fig. 3i** and **Supplementary Table 4**). Interestingly, ACAT1 harbors a highly KB02-reactive cysteine (C126)^14, 15^, which suggests that the direct engagement by KB02- containing compounds like KB02-JQ1 could lead to degradation of this protein. Supporting a functional role for DCAF16 in KB02-JQ1-induced BRD4 degradation, we found that the HMW form of DCAF16 co-immunoprecipitated with BRD4 in a KB02-JQ1-dependent manner (**Fig. 3j**) and that BRD4 degradation was substantially blocked in DCAF16-/- cells (**Fig. 3k**).

In summary, we have used a chemical proteomic strategy to discover that the E3 ligase subunit DCAF16 supports targeted protein degradation when engaged by electrophilic PROTACs. DCAF16 may offer advantages as a new addition to the emerging subset of E3 ligases that promote ligand-induced protein degradation. First, our data suggest that DCAF16 exclusively promotes the degradation of nuclear proteins, which may offer a way to improve the specificity of ligand-induced protein degradation by, for instance, avoiding cytosolic proteins that engage a bifunctional compound. Second, DCAF16 appears capable of supporting ligand-induced protein degradation at very low fractional engagement, which we attribute, at least in part, to covalent interaction with electrophilic PROTACs. By converting a sub-stoichiometric portion of DCAF16 to a ‘neo-E3 ligase’, electrophilic PROTACs may support the degradation of target proteins while minimally perturbing endogenous substrates of DCAF16, which could presumably be degraded by the still predominantly unmodified fraction of this E3 ligase in cells. Our findings also underscore the value of broadly reactive scout fragments as tools to discover ligandable and functional sites on poorly characterized proteins like DCAF16. We should note, however, that scout fragments, even when active as electrophilic PROTACs at sub-μM concentrations, still modify several other proteins in cells, and the optimization of ligands for DCAF16-dependent protein degradation therefore represents an important future objective. Here, structural information on DCAF16, in particular, adducted to covalent PROTACs, would be valuable, as our chemical proteomic data point to a site(s) of modification that contains several clustered cysteines. We finally wonder whether this feature indicates the potential for endogenous electrophiles and/or oxidative stress to modify DCAF16 and shape its substrate scope *in vivo*.

## Acknowledgements

This work was supported by the NIH (CA087660 (B.F.C.), CA231991 (B.F.C.), CA212467 (M.M.D.), CA211526 (V.M.C.)) and the Damon-Runyon Cancer Research Foundation (X.Z. DRG-2341-18).

## Author contributions

X.Z. and B.F.C. conceived of the research and wrote the paper. X.Z. developed methods, performed experiments, and analyzed data. X.Z. and M.M.D. analyzed chemical proteomic data. X.Z. designed and synthesized KB02-SLF, KB02-PEG0-SLF, KB02-PEG4-SLF, KB03-SLF, KB05-SLF and C-KB02-SLF. V.M.C. designed and synthesized KB02-JQ1. T.G.W. designed and synthesized lenalidomide-SLF. V.M.C. characterized all the compounds.

---

**Supplementary Fig. 1.**
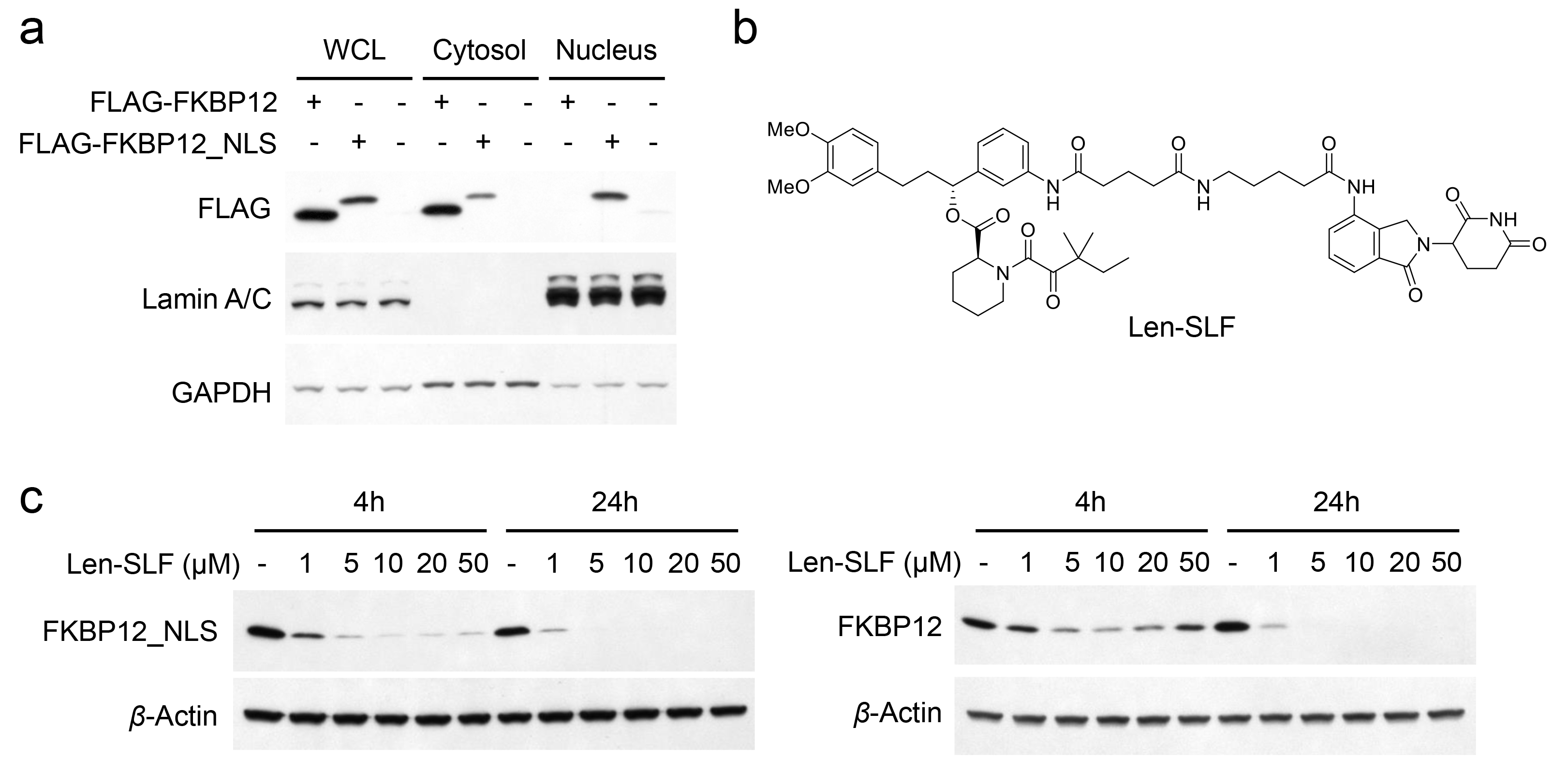
Stable expression of FLAG-FKBP12 and FLAG-FKBP12_NLS in HEK293T cells and characterization of a lenalidomide-SLF PROTAC. **a**, Subcellular fractionation followed by western blotting of stably expressed FLAG-FKBP12 and FLAG-FKBP12_NLS in HEK293T cells. The result is a representative of two independent experiments. **b**, Structure of lenalidomide-SLF. **c**, Concentration-dependent degradation of stably expressed FLAG-FKBP12 and FLAG-FKBP12_NLS in HEK293T cells following treatment with lenalidomide-SLF for 4 or 24 h.

**Supplementary Fig. 2.**
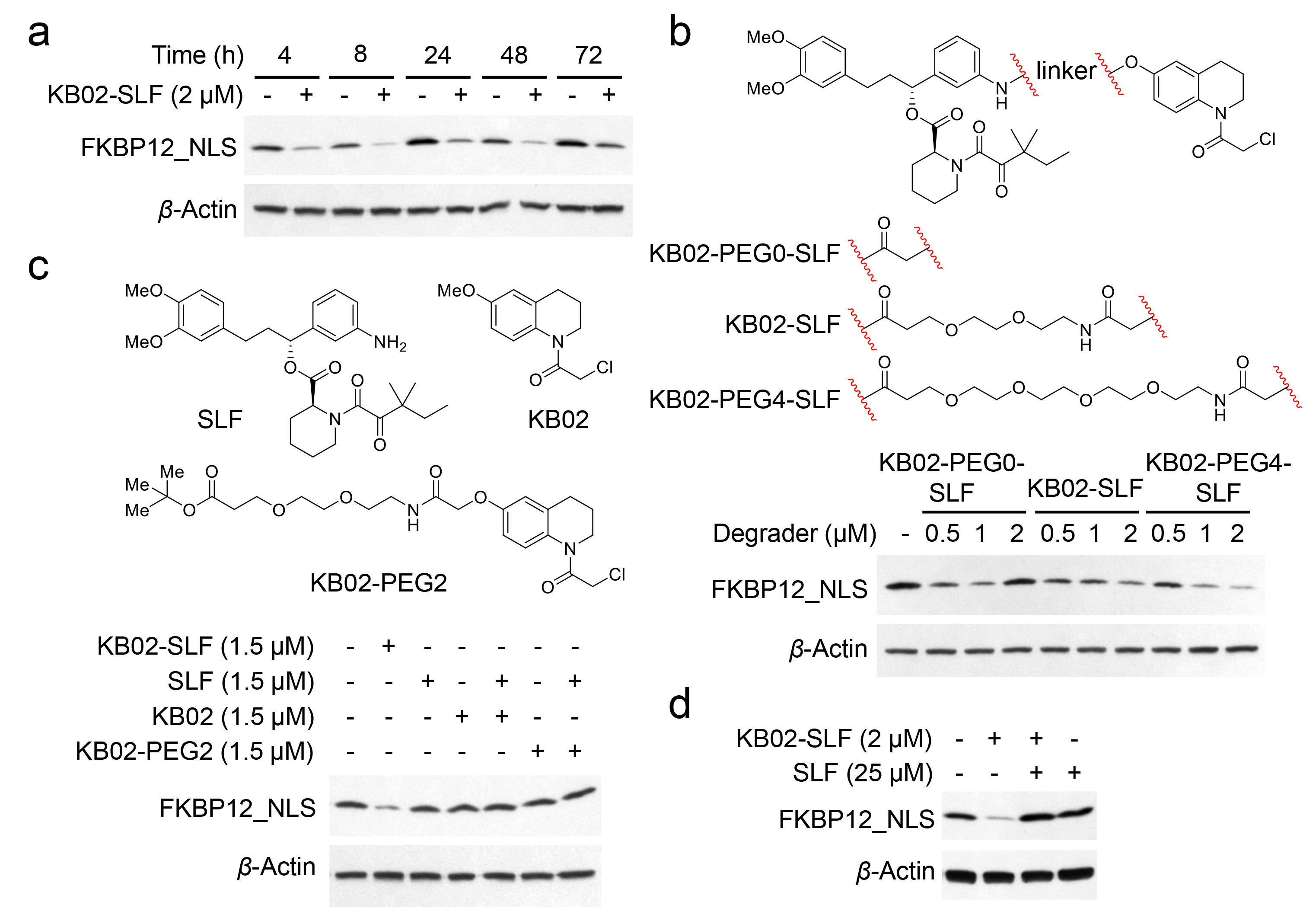
Characterization of KB02-SLF PROTACs. **a**, Time-dependent degradation of FLAG-FKBP12_NLS in HEK293T cells treated with KB02-SLF (2 μM). The result is a representative of two independent experiments. **b**, Top, structures of KB02-PEG0-SLF, KB02-SLF, and KB02-PEG4-SLF. Bottom, western blot of FLAG-FKBP12_NLS following treatment of HEK293T cells with KB02-PEG0-SLF, KB02-SLF, or KB02-PEG4-SLF (0.5-2 μM, 8 h). The result is a representative of three independent experiments. **c**, Western blot of FLAG-FKBP12_NLS in HEK293T cells following treatment of HEK293T cells with KB02, KB02-PEG2, SLF, or the combination of SLF and KB02 or KB02-PEG2 (1.5 μM, 8 h). **d**, Degradation of FLAG-FKBP12_NLS by KB02-SLF (2 μM, 8 h) is blocked by excess SLF (25 μM). The result is a representative of three independent experiments.

**Supplementary Fig. 3.**
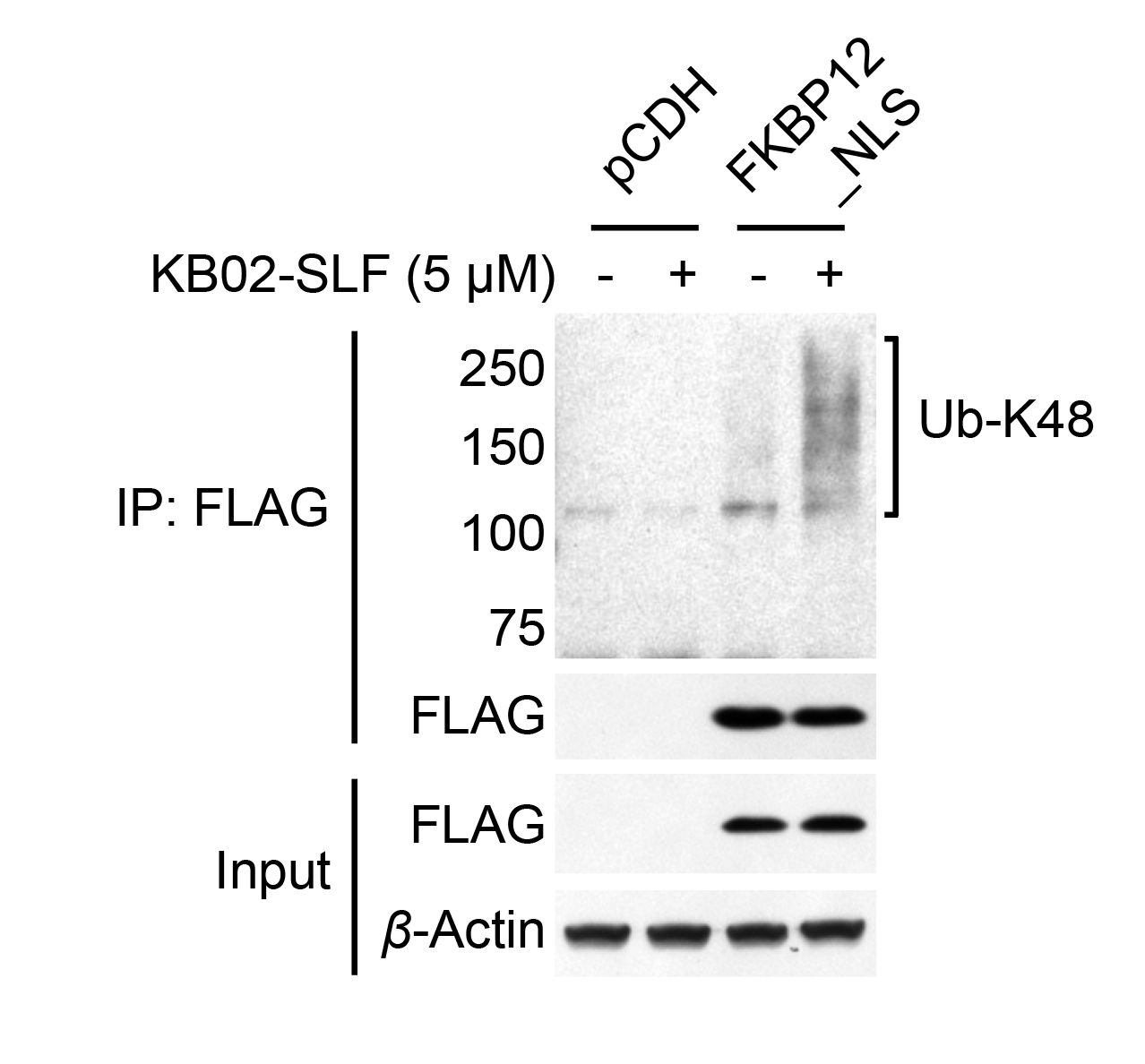
KB02-SLF induces K48-linked polyubiquitination on FLAG-FKBP12_NLS. HEK293T cells stably expressing FLAG-FKBP12_NLS were treated with KB02- SLF (5 μM) and MG132 (10 μM) for 2 h, and FLAG-FKBP12_NLS was then immunoprecipitated and analyzed by western blotting for K48-linked ubiquitination. The result is a representative of two independent experiments.

**Supplementary Fig. 4.**
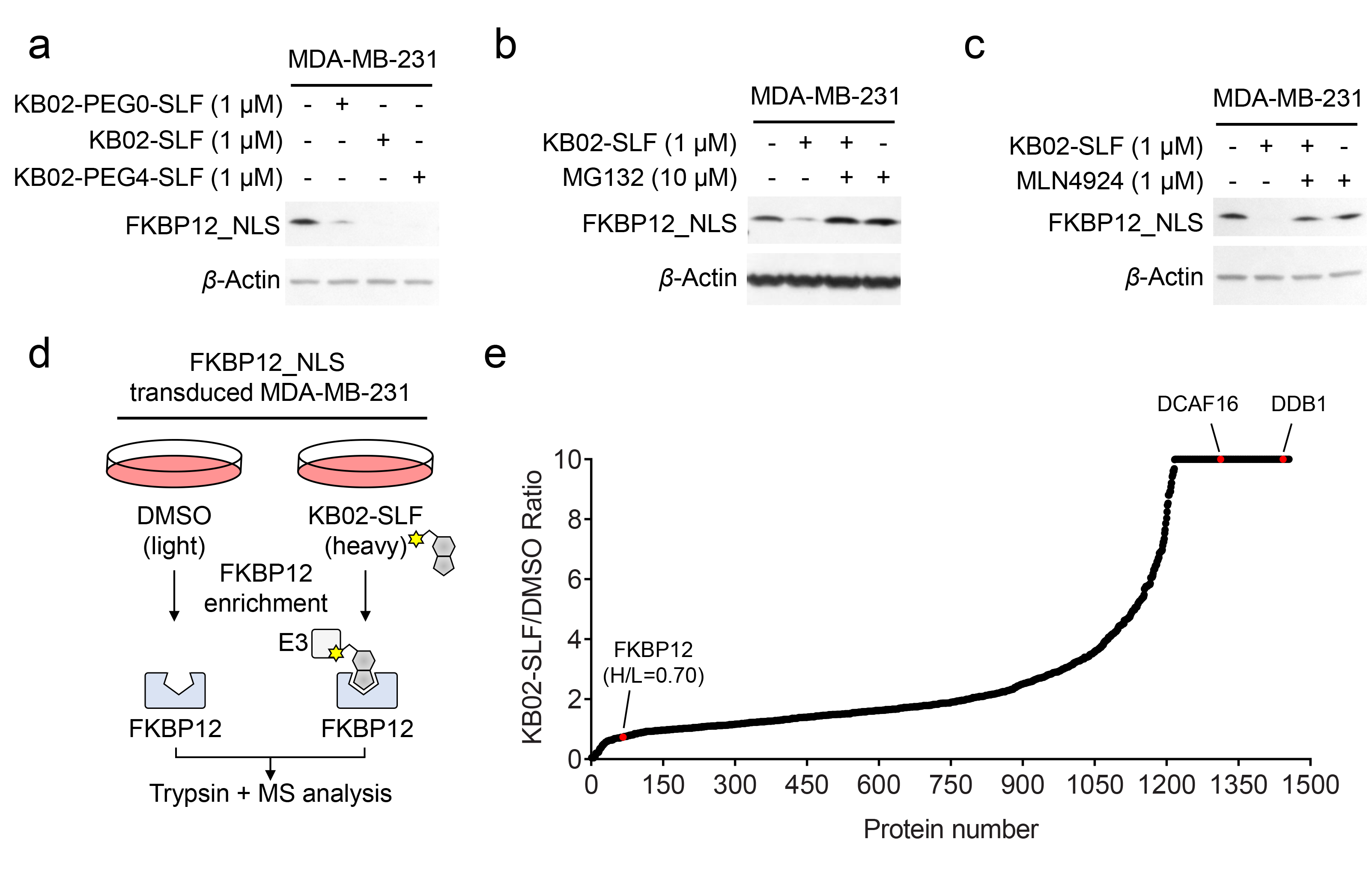
KB02-SLF degrades FLAG-FKBP12_NLS in MDA-MB-231 cells. **a**, Western blot of stably expressed FLAG-FKBP12_NLS in MDA-MB-231 cells following treatment with KB02-PEG0-SLF, KB02-SLF or KB02-PEG4-SLF (1 μM, 8 h). The result is a representative of two independent experiments. **b**, **c**, KB02-SLF-mediated degradation of FLAG-FKBP12_NLS is blocked by the proteasome inhibitor MG132 (**b**) and the neddylation inhibitor MLN4924 (**c**) in MDA-MB-231 cells. MDA-MB-231 cells stably expressing FLAG-FKBP12_NLS were co-treated with KB02-SLF (1 μM) and MG132 (10 μM) or MLN4924 (1 μM) for 8 h. The result is a representative of three independent experiments. **d**, Schematic for identifying KB02-SLF-recruited E3 ubiquitin ligases by anti-FLAG affinity enrichment coupled to mass spectrometry (MS)-based proteomics. Light and heavy amino acid-labeled MDA-MB-231 cells stably expressing FLAG-FKBP12_NLS were treated with DMSO or KB02-SLF (10 μM), respectively, for 2 h in the presence of MG132 (10 μM). Light and heavy cells were then lysed, subject to anti-FLAG immunoprecipitation, and the affinity-enriched proteins combined, digested with trypsin, and analyzed by LC-MS/MS. **e**, SILAC heavy/light (H/L) ratio values of proteins identified in anti-FLAG affinity enrichment experiments (outlined in part **d**), where a high ratio indicates proteins selectively enriched from cells treated with KB02-SLF.

**Supplementary Fig. 5.**
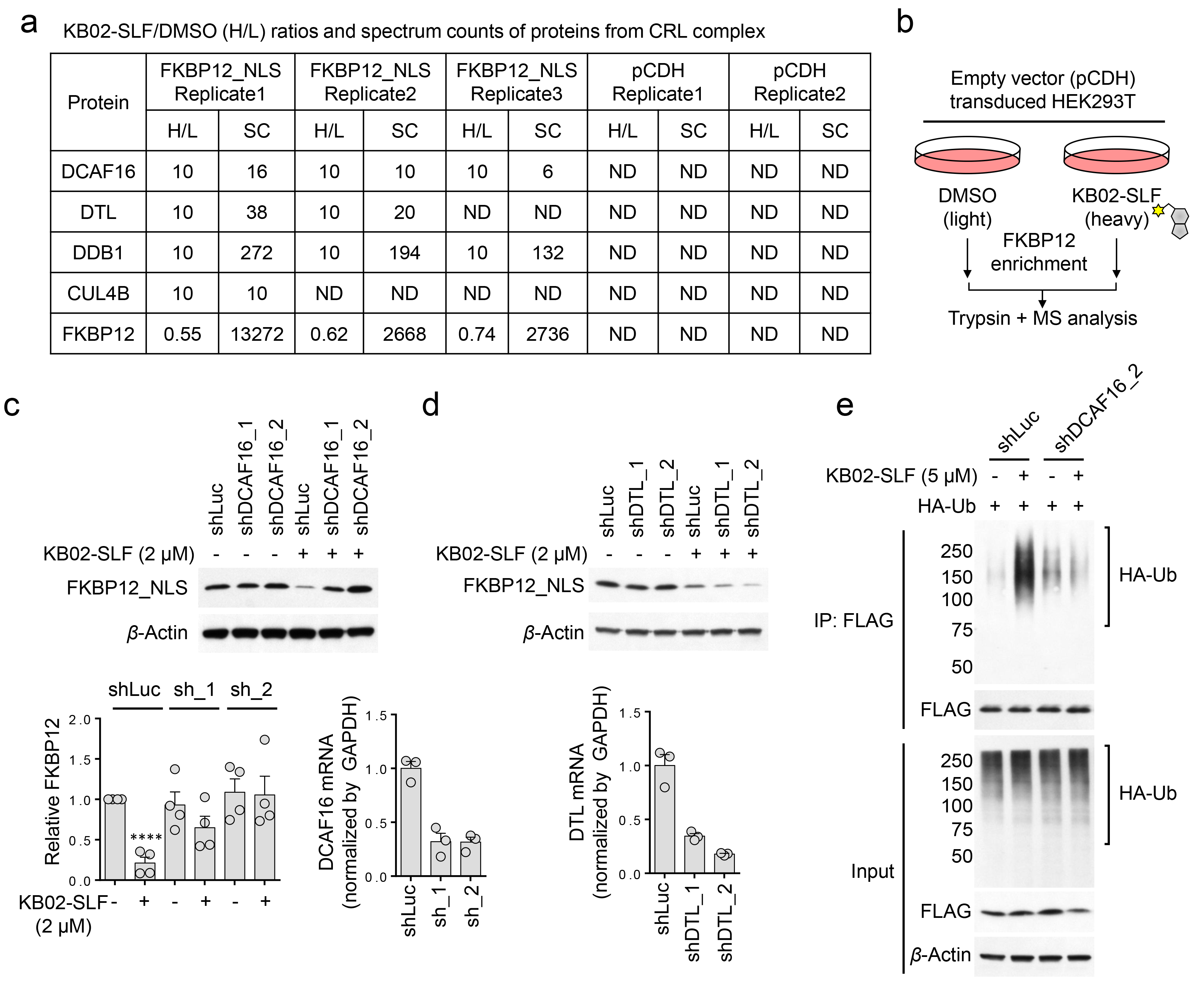
Characterization of KB02-SLF-recruited E3 ubiquitin ligase(s) in HEK293T cells by anti-FLAG affinity enrichment coupled to mass spectrometry (MS)-based proteomics. **a**, SILAC H/L ratio and spectral count (SC) values for E3 ligase proteins and FKBP12 enriched by anti-FLAG immunoprecipitation from HEK239T cells stably expressing FLAG-FKBP12_NLS, but not enriched from control HEK293T cells expressing empty pCDH vector. **b**, Schematic for a control affinity-enrichment-MS-based proteomics experiment, where light and heavy amino acid-labeled HEK293T cells stably expressing pCDH empty vector were treated for 2 h with DMSO or KB02-SLF (10 μM), respectively in the presence of 10 μM MG132 (10 μM). Light and heavy cells were then lysed, subject to anti-FLAG immunoprecipitation, and proteins bound to anti-FLAG beads combined, digested with trypsin, and analyzed by LC-MS/MS. **c**, Western blot of stably expressed FLAG-FKBP12_NLS in HEK293T cells transiently transduced with shRNAs targeting DCAF16 (sh_1 and sh_2) or a control shRNA (shLuc) followed by treatment with KB02-SLF (2 μM, 8 h). Bottom left, quantification of the relative FKBP12 protein content, with DMSO-treated cells expressing shLuc set to a value of 1. Data represent mean values ± SEM for 4 biological replicates. Statistical significance was calculated with unpaired two-tailed Student’s t-tests comparing DMSO- to KB02-SLF-treated samples. \*\*\*\**P* < 0.0001. Bottom right, DCAF16 mRNA as measured by qPCR. Data represent mean values ± SEM for 3 biological replicates. **d**, Top, western blot of stably expressed FLAG-FKBP12_NLS in HEK293T cells transiently transduced with shRNAs targeting DTL (shDTL) or a control (shLuc) following treatment with KB02-SLF (2 μM, 8 h). Bottom, DTL mRNA was measured by qPCR. Data represent mean values ± SEM for 3 biological replicates. **e**, shRNA-mediated DCAF16 knockdown attenuates KB02-SLF-dependent polyubiquitination of FLAG-FKBP12_NLS. HEK293T cells stably expressing FLAG-FKBP12_NLS were transiently transduced with an shRNA targeting DCAF16 (shDCAF16) or a control shRNA (shLuc) and treated with KB02-SLF (5 μM) and MG132 (10 μM) for 2 h. The result is a representative of two independent experiments.

**Supplementary Fig. 6.**
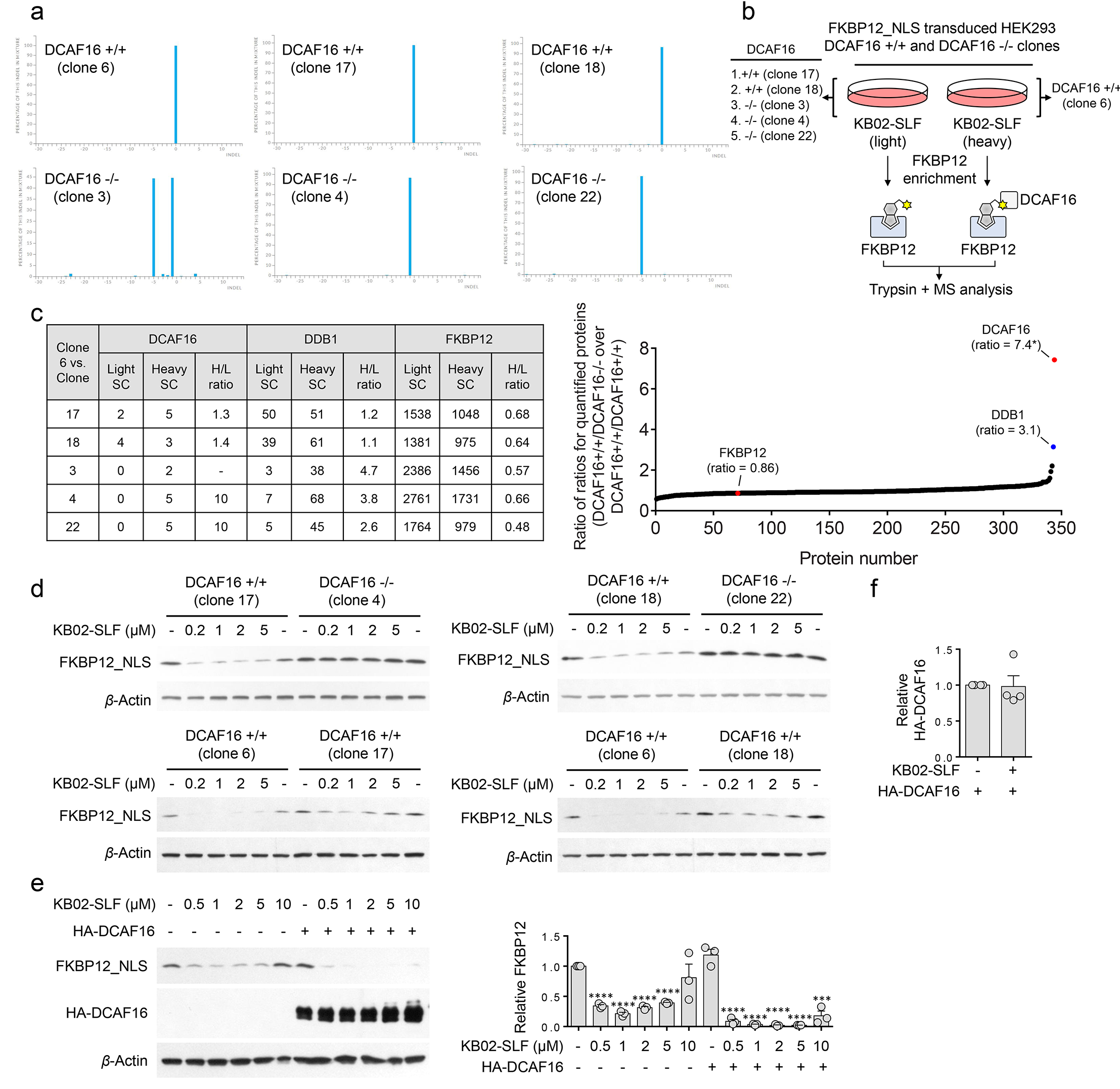
Generation of DCAF16-/- HEK293 cells using CRISPR/Cas9 and characterization of KB02-SLF-mediated degradation of FLAG-FKBP12_NLS in these cells. **a**, Indel analysis of three DCAF16+/+ clones (clones 6, 17, and 18) and three DCAF16-/- clones (clones 3, 4, and 22) in HEK293 cells. **b**, Schematic for measuring DCAF16 content in DCAF16+/+ and DCAF16-/- clones by anti-FLAG affinity enrichment coupled to MS-based proteomics of KB02-SLF-treated HEK293 cells stably expressing FLAG-FKBP12_NLS. Light and heavy amino acid-labeled cells from clones shown in the schematic were treated with DMSO or KB02-SLF (10 μM), respectively, for 2 h in the presence of MG132 (10 μM). Light and heavy cells were then lysed, subject to anti-FLAG immunoprecipitation, and the affinity-enriched proteins combined, digested with trypsin, and analyzed by LC-MS/MS. **c**, DCAF16 content from indicated DCAF16+/+ and DCAF16-/- clones analyzed as described in part **b**. *Note that a maximum H/L value of 10 was assigned to DCAF16 in DCAF16+/+/DCAF16-/- comparisons, which results in a calculated ratio of 7.4 for DCAF16 in the waterfall plot (right) showing ratio values for proteins in DCAF16+/+/DCAF16-/- versus DCAF16+/+/DCAF16+/+ comparisons. **d**, Concentration-dependent degradation of stably expressed FLAG-FKBP12_NLS in DCAF16+/+ and DCAF16-/- HEK293 cells treated with KB02-SLF (0.2-5 μM, 8 h). The result is a representative of two independent experiments. **e**, Concentration-dependent degradation of FLAG-FKBP12_NLS by KB02-SLF in HEK293T cells with or without HA-DCAF16 transfection (8 h treatment with indicated concentrations of KB02-SLF). Bar graph (right) represents quantification of the relative FKBP12 protein, with DMSO-treated cells lacking HA-DCAF16 transfection set to a value of 1. Data represent mean values ± SEM for 3 biological replicates. Statistical significance was calculated with unpaired two-tailed Student’s t-tests comparing DMSO- to KB02-SLF-treated samples. \*\*\**P* < 0.001; \*\*\*\**P* < 0.0001. **f**, Quantification of the relative HA-DCAF16 content in **Fig. 3c**, where the HA-DCAF16 level with DMSO-treated cells set to a value of 1. Data represent mean values ± SEM for 4 biological replicates (two from DCAF16-/- clone 3, two from DCAF16-/- clone 4). Statistical significance was calculated with unpaired two-tailed Student’s t-tests comparing DMSO- to KB02-SLF-treated samples.

**Supplementary Fig. 7.**
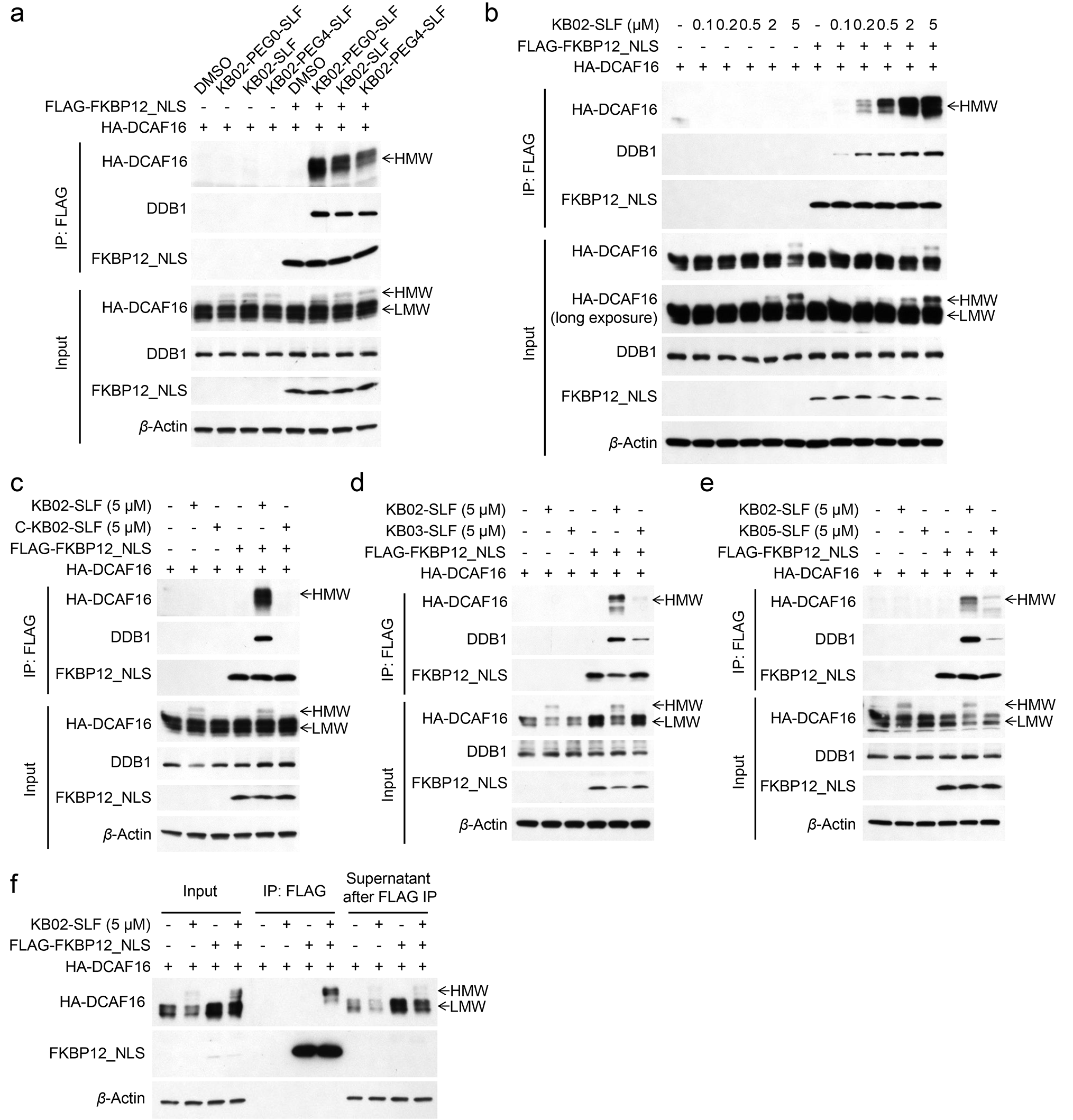
KB02-SLF mediates a ternary complex interaction between FLAG-FKBP12_NLS and HA-DCAF16. **a**, The higher molecular weight (HMW) form of FLAG-FKBP12_NLS co-immunoprecipitated with HA-DCAF16 in the presence of KB02-PEG0-SLF, KB02-SLF or KB02-PEG4-SLF. HEK293T cells stably expressing FLAG-FKBP12_NLS were transfected with HA-DCAF16 for 24 h and then treated with KB02-PEG0-SLF, KB02-SLF or KB02-PEG4-SLF (5 μM) in the presence of MG132 (10 μM) for 2 h. The result is a representative of two independent experiments. **b**, Concentration-dependent co-immunoprecipitation of the HMW form of FLAG-FKBP12_NLS with HA-DCAF16 in HEK293T cells treated with KB02-SLF in the presence of MG132 (10 μM) for 2 h. The result is a representative of two independent experiments. **c-e**, FLAG-FKBP12_NLS co-immunoprecipitated with HA-DCAF16 in the presence of KB02-SLF, but not C-KB02-SLF, KB03-SLF, or KB05-SLF. HEK293T cells stably expressing FLAG-FKBP12_NLS were transfected with HA-DCAF16 for 24 h and the treated with KB02-SLF, C-KB02-SLF, KB03-SLF or KB05-SLF (5 μM) in the presence of MG132 (10 μM) for 2 h. The result is a representative of two independent experiments. **f**, The higher molecular weight (HMW) form of FLAG-FKBP12_NLS co-immunoprecipitated with HA-DCAF16 in the presence of KB02-SLF. HEK293T cells stably expressing pCDH empty vector or FLAG-FKBP12_NLS were transfected with HA-DCAF16 for 24 h and then treated with DMSO or KB02-SLF (5 μM) in the presence of MG132 (10 μM) for 2 h. The result is a representative of two independent experiments.

**Supplementary Fig. 8.**
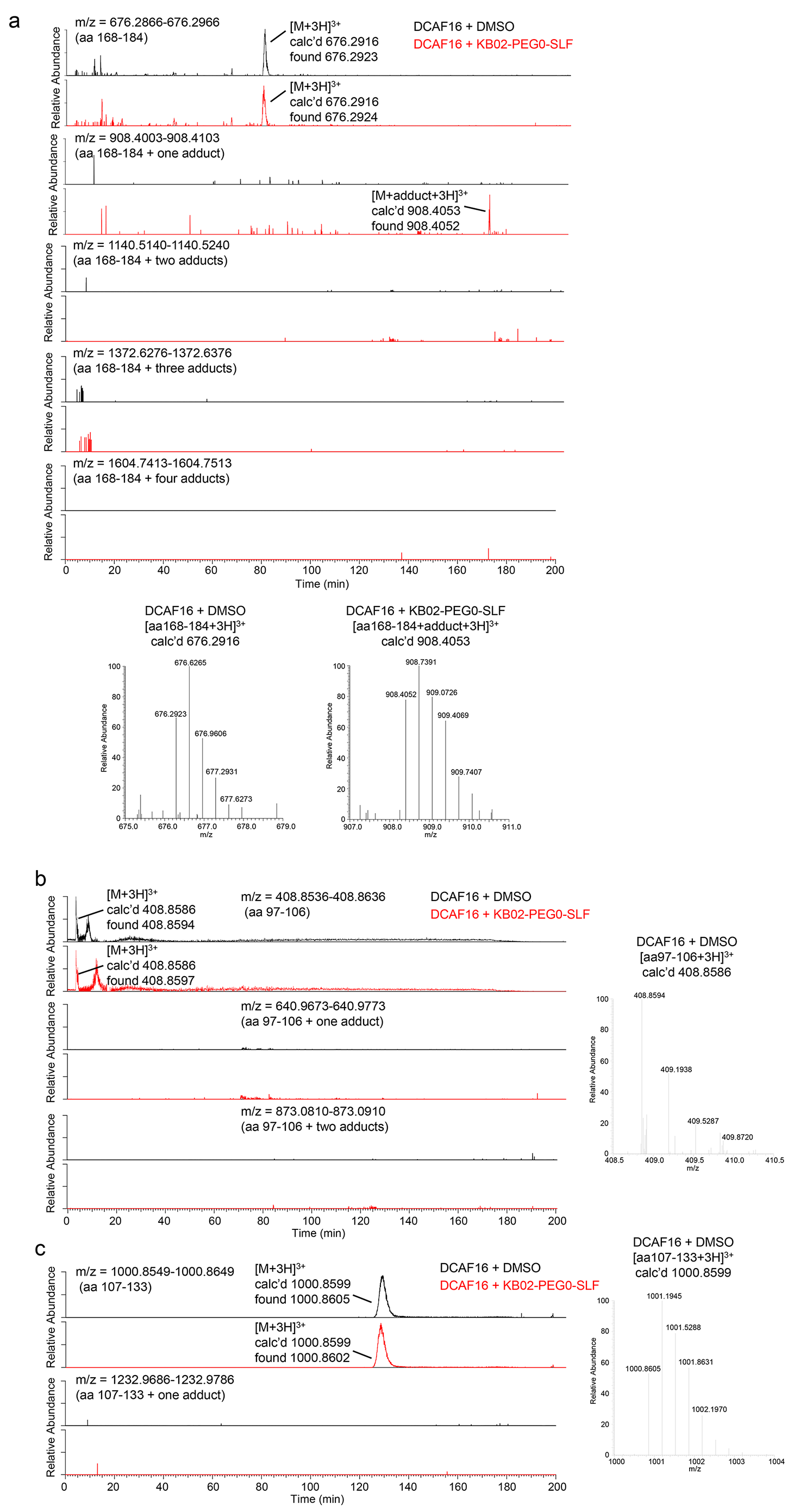
MS analysis of KB02-PEG0-SLF-modified cysteine(s) in DCAF16. **a**, Top: extracted ion chromatograms (EICs) for DCAF16 tryptic peptide (amino acids 168-184) with or without KB02-PEG0-SLF modification. Bottom: MS1 spectra of triply charged DCAF16 tryptic peptide (amino acids 168-184) and KB02-PEG0-SLF-modified, triply charged DCAF16 tryptic peptide (amino acids 168-184). **b**, Left: extracted ion chromatograms (EICs) for DCAF16 tryptic peptide (amino acids 97-106) with or without the predicted KB02-PEG0-SLF modification. Right: MS1 spectrum of triply charged DCAF16 tryptic peptide (amino acids 97-106). **c**, Left: extracted ion chromatograms (EICs) for DCAF16 tryptic peptide (amino acids 107-133) with or without the predicted KB02-PEG0-SLF modification. Right: MS1 spectrum of triply charged DCAF16 tryptic peptide (amino acids 107-133).

**Supplementary Fig. 9.**
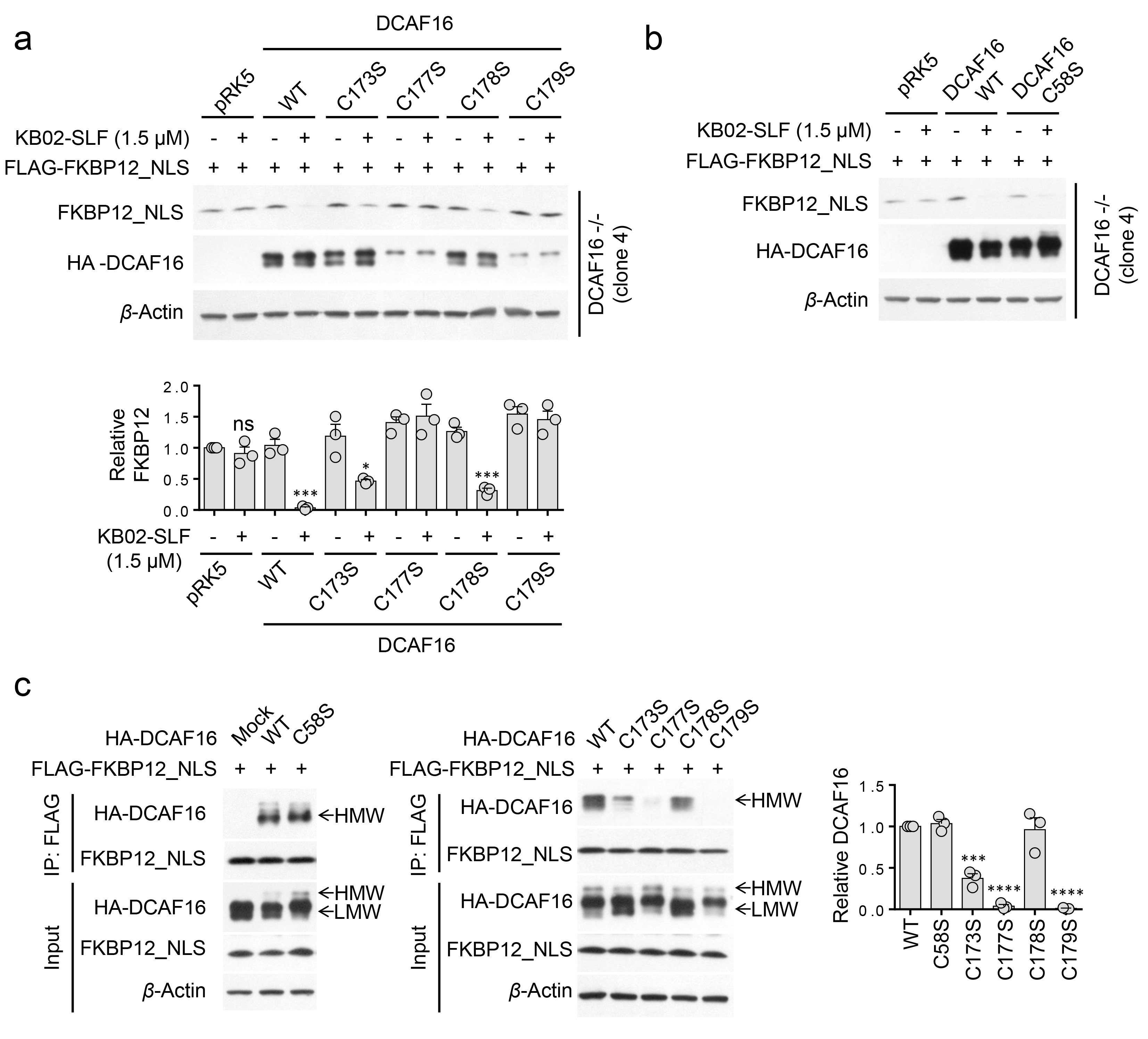
Analysis by site-directed mutagenesis of the contributions of cysteines on the KB02-SLF-modified DCAF16 peptide (amino acids 168-184) to KB02- SLF-mediated degradation of FLAG-FKBP12_NLS. **a**, Western blot of FLAG-FKBP12_NLS in DCAF16-/- HEK293 cells expressing WT or C173S, C177S, C178S, or C179S mutants of HA-DCAF16 (or empty pRK5 vector control) following treatment with KB02-SLF (1.5 μM, 8 h). Bar graph (bottom) represents quantification of the relative FKBP12 protein content, with DMSO-treated cells set to a value of 1. Data represent mean values ± SEM for 3 biological replicates. Statistical significance was calculated with unpaired two-tailed Student’s t-tests comparing DMSO- to KB02-SLF-treated samples. \*\*\**P* < 0.001. **b**, Western blot of FLAG-FKBP12_NLS in DCAF16-/- HEK293 cells expressing WT or C58S mutant of HA-DCAF16 (or empty pRK5 vector control) following treatment with KB02-SLF (1.5 μM, 8 h). The result is a representative of three independent experiments. **c**, KB02-SLF-mediated interaction between the HMW form of WT or mutant HA-DCAF16 and FLAG-FKBP12_NLS as determined by co-immunoprecipitation. Bar graph represents quantification of the relative HA-DCAF16 protein content, with DMSO-treated cells set to a value of 1. Data represent mean values ± SEM for 3 biological replicates. Statistical significance was calculated with unpaired two-tailed Student’s t-tests comparing DCAF16 mutant to DCAF16 WT. \*\*\**P* < 0.001; \*\*\*\**P* < 0.0001.

**Supplementary Fig. 10.**
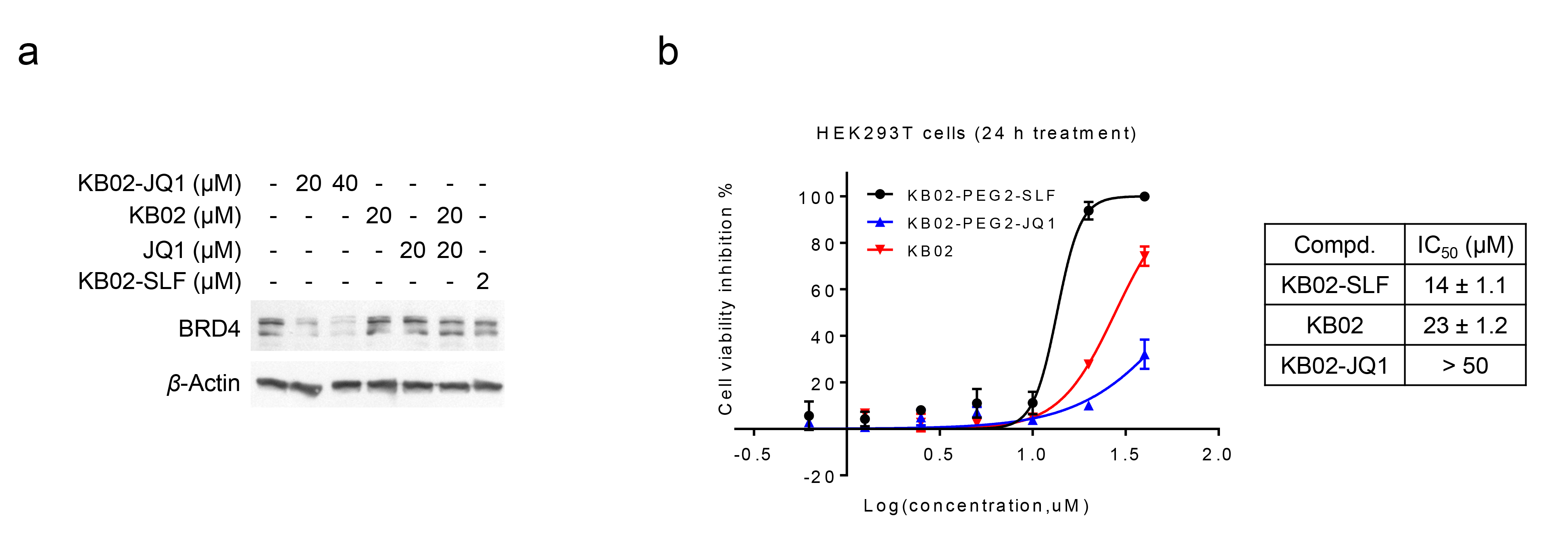
**a**, Western blot of BRD4 in HEK293T cells following treatment with the indicated concentrations of KB02-JQ1, KB02, JQ1, KB02-SLF, or the combination of KB02 and JQ1 (24 h). **b**, Cell viability of HEK293T cells treated with KB02-SLF, KB02, or KB02-JQ1 for 24 h. Data represent mean values ± SEM for 3 biological replicates.

**Supplementary Table 1**. isoTOP-ABPP data for HEK293T cells treated with KB02-SLF (10 μM, 2 h) or DMSO.

See accompanying Excel file.

**Supplementary Table 2.** Complete proteomic data for light and heavy amino acid-labeled HEK293T cells stably expressing FLAG-FKBP12_NLS that were treated with DMSO or KB02- SLF (10 μM), respectively, for 2 h in the presence of MG132 (10 μM), lysed, subject to anti-FLAG immunoprecipitation, and the affinity-enriched proteins combined, digested with trypsin, and analyzed by LC-MS/MS. Related to **Fig. 2d, e**.

See accompanying Excel file.

**Supplementary Table 3.** Complete proteomic data for comparison of DCAF16+/+ and DCAF16-/- clones by anti-FLAG affinity enrichment coupled to mass spectrometry-based proteomics of KB02-SLF-treated HEK293 cells stably expressing FLAG-FKBP12_NLS. Related to **Supplementary Fig. 6b, c**.

See accompanying Excel file.

**Supplementary Table 4.** Proteomics data for fold-change in protein abundance between heavy- and light-isotopically labeled HEK293T cells treated with KB02-JQ1 (20 μM)/DMSO or DMSO/DMSO, respectively, for 24 h. Related to **Fig. 3i**.

See accompanying Excel file.

